# Accurate brain age prediction with lightweight deep neural networks

**DOI:** 10.1101/2019.12.17.879346

**Authors:** Han Peng, Weikang Gong, Christian F. Beckmann, Andrea Vedaldi, Stephen M. Smith

**Author notes:** These authors contributed equally (Han Peng, Weikang Gong). Corresponding to Han Peng.

## Abstract

Deep learning has huge potential for accurate disease prediction with neuroimaging data, but the prediction performance is often limited by training-dataset size and computing memory requirements. To address this, we propose a deep convolutional neural network model, Simple Fully Convolutional Network (SFCN), for accurate prediction of brain age using T1-weighted structural MRI data. Compared with other popular deep network architectures, SFCN has fewer parameters, so is more compatible with small dataset size and 3D volume data. The network architecture was combined with several techniques for boosting performance, including data augmentation, pre-training, model regularization, model ensemble and prediction bias correction. We compared our overall SFCN approach with several widely-used machine learning models. It achieved state-of-the-art performance in UK Biobank data (N = 14,503), with mean absolute error (MAE) = 2.14y in brain age prediction and 99.5% in sex classification. SFCN also won (both parts of) the 2019 Predictive Analysis Challenge for brain age prediction, involving 79 competing teams (N = 2,638, MAE = 2.90y). We describe here the details of our approach, and its optimisation and validation. Our approach can easily be generalised to other tasks using different image modalities, and is released on GitHub.

**Highlights:**

- A lightweight deep learning model, Simple Fully Convolutional Network (SFCN), is presented, achieving state-of-the-art brain age prediction and sex classification performance in UK Biobank MRI brain imaging data.
- Even with limited number of training subjects (e.g., 50), SFCN performs better than widely-used regression models.
- A semi-multimodal ensemble strategy is proposed and achieved first place in the PAC 2019 brain age prediction challenge.
- Linear regression can remove brain age prediction bias (even on unlabelled data) while maintaining state-of-the-art performance.

## 1 Introduction

The emergence of machine learning techniques has made automatic disease prediction from medical imaging data possible. The recent development of deep learning pushes prediction accuracy beyond human performance in some scenarios, and is able to assist clinical diagnosis/treatment decisions (De Fauw et al., 2018; Kohl et al., 2018; LeCun et al., 2015). In neuroimaging, deep learning has had successes in several applications in predictive and diagnostic analysis, such as brain age prediction and modelling (Cole et al., 2017; Kawahara et al., 2017), sex classification (Arslan et al., 2018), disease prediction (Baumgartner et al., 2018; Korolev et al., 2017; Liu et al., 2018) and brain lesion segmentation (Kamnitsas et al., 2017), yet still faces several challenges. For example, 3D neuroimaging data requires much more GPU memory than most 2D images, meaning that models successful in 2D data (e.g., ImageNet classification (Krizhevsky et al., 2012; Simonyan and Zisserman, 2014)) are infeasible in the 3D scenario. There are several researches mitigating this issue (e.g.) by downsampling the input (Korolev et al., 2017), taking patches (Kamnitsas et al., 2017; Liu et al., 2018) or 2D slices (Bashyam et al., 2020; Lin et al., 2018) as input instead of the 3D full brain, or using a reversible architecture (Brügger et al., 2019), yet involving trade-offs between the GPU memory restriction and the information/performance loss. Further, deep networks usually require a large sample size for model fitting, but neuroimaging datasets often have relatively few samples compared to existing million-sample natural image datasets (Raghu et al., 2019; Russakovsky et al., 2015), which could limit the ability to learn image features effectively, and result in overfitting problems. New model architecture design is needed to address these challenges for neuroimaging applications.

Predicting chronological age based on structural brain magnetic resonance imaging (MRI) data shares common challenges with many other neuroimaging applications, and can be used to develop and test deep learning algorithms. It also receives attention for its potential clinical and biological relevance (Ashburner, 2007; Brown et al., 2012; Cole et al., 2018; Cole and Franke, 2017; Davatzikos et al., 2009; Franke et al., 2014, 2010; Franke and Gaser, 2019; Habes et al., 2016; Kaufmann et al., 2019; Neeb et al., 2006). The predicted age can be considered to be the “brain age”, because it is derived purely from the brain imaging data. After estimating brain age, a further quantity of interest is the difference between the predicted age (brain age) and the actual age, sometimes referred to as the brain-age delta. Positive delta implies that a subject’s brain looks older than their actual age, i.e., they are experiencing accelerated aging. For example, existing studies have observed that the brain-age delta is an effective biomarker that shows differences between different clinical groups (Kaufmann et al., 2019) and is predictive for mortality (Cole et al., 2018). Achieving accurate brain age prediction is an essential pre-requisite for optimising brain-age delta as a biomarker. To reach this goal, many studies have used different models, such as regularized linear regression, support vector machines and Gaussian process regression, for brain age prediction (Franke and Gaser, 2019). Some studies have used deep learning methods (Cole et al., 2017; Feng et al., 2019; Kolbeinsson et al., 2019). However, challenges exist for further improvement of prediction accuracy, especially on small datasets, and some studies have shown that deep learning performs no better than simpler machine learning models in typical neuroimaging datasets (He et al., 2020; Schulz et al., 2019). It has not yet, for example, been clearly established whether more complex deep learning models perform better than simpler models (for the task of brain age prediction using structural MRI data). In addition, predicted brain age is often systematically biased towards the group mean value, resulting in a correlation between the delta and the chronological age, which weakens the validity of the delta as a biomarker (Smith et al., 2019). Therefore, it is both methodologically interesting and scientifically important to develop unbiased high-performance deep learning strategies for brain age prediction.

In this paper, a lightweight deep learning architecture, Simple Fully Convolutional Network (SFCN), is presented for brain age prediction. Its architecture is based on the fully convolutional network (FCN) (Long et al., 2015) and the VGG net (Simonyan and Zisserman, 2014) and takes 3D minimally-preprocessed T1 brain images (and/or preprocessed segmentation outputs) as input. The successful CNN architecture VGG net and its variant with BatchNorm layers (Ioffe and Szegedy, 2015) provide a deep architecture consisting of a sequence of basic blocks: (Conv-BatchNorm-Activation)xN-Pooling. To reduce memory requirements, SFCN keeps only one conv-layer before each MaxPool layer. In addition, we remove all the fully-connected layers, which not only greatly reduces the number of parameters, but also provides a fully convolutional architecture that is versatile for accommodating different input sizes (Long et al., 2015). Using proper data augmentation and regularization techniques, the model achieved state-of-the-art mean absolute error (MAE) of 2.14 years in the UK Biobank dataset (14,503 subjects, of which 12,949 are used for training). This model performs better than several widely-used machine learning models in the literature. In addition, we propose a model ensemble strategy that averages the outputs of deep learning models based on different kinds of preprocessing applied to the T1 data, namely, white matter segmentation, grey matter segmentation, linearly-registered raw T1 and nonlinearly registered raw T1; this further boosts the accuracy of brain age prediction. Finally, we extended the bias correction techniques proposed by (Smith et al., 2019) to greatly reduce the correlation between the brain-age delta and the chronological age, with very little compromise of performance, even when the true ages of test (validation) subjects are unknown. The ensemble SFCN came first in the Predictive Analysis Challenge 2019 in brain age prediction (MAE = 2.90 years) among the 79 participating teams^1^. With bias correction, our model achieved an MAE of 2.95 years, thereby ranking first also in the other part of the competition (most accurate age prediction while minimising bias). The trained model is available in the GitHub repository: https://github.com/ha-ha-ha-han/UKBiobank_deep_pretrain/

Our main contributions in this paper are:

1. We propose a novel lightweight 3D CNN architecture, Simple Fully Convolutional Network, which performances better than deeper CNNs and is even more data efficient than simpler linear models in brain age and sex prediction.
2. We demonstrate that combining complementary information from different preprocessing pipelines improve age prediction accuracy (even by simply averaging the outputs using different modalities).
3. We propose a novel bias correction method for brain age prediction which can be applied to new *unknown-label* test datasets.

## 2 Methods

### 2.1 Model: Simple Fully Convolutional Network (SFCN)

We use a convolutional neural network (CNN) architecture to estimate brain age using 3D T1 images. The architecture is based on VGGNet (Simonyan and Zisserman, 2014) and uses a fully convolutional structure (Long et al., 2015), but we keep the number of layers as small as possible to reduce the number of parameters to about 3 million, and therefore to reduce computational complexity and memory cost. We name this model structure “Simple Fully Convolutional Neural Network” (SFCN) to reflect its simplicity.

The model consists of seven blocks, as shown in Figure 1. Each of the first five blocks contains a 3-by-3-by-3 3D convolutional layer, a batch normalisation layer (Ioffe and Szegedy, 2015), a max pooling layer and a ReLU activation layer (LeCun et al., 2015). The 1mm-input-resolution 160×192×160 3D input image (with little or no brain tissue loss) goes through each block sequentially, with its feature map generated and spatial dimension reduced to 5×6×5 after the fifth block. The sixth block contains a 1×1×1 3D convolutional layer, a batch normalisation layer and a ReLU activation layer. The seventh block contains an average pooling layer, a dropout layer (only used for training, with 50% random dropout rate) (Srivastava et al., 2014), a fully connected layer and a softmax output layer. The channel numbers used in each convolution layer are [32, 64, 128, 256, 256, 64, 40]. The output layer contains 40 digits that represent the predicted probability that the subject’s age falls into a one-year age interval between 42 to 82 (for UK Biobank) or a two-year age interval between 14 to 94 (for PAC 2019). A weighted average of each age bin is calculated to make the final prediction:

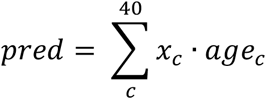

*x*_c_ stands for the probability predicted for the *c*^th^ age bin and *age*_*c*_ stands for the bin centre for the age interval.

**Figure 1.**
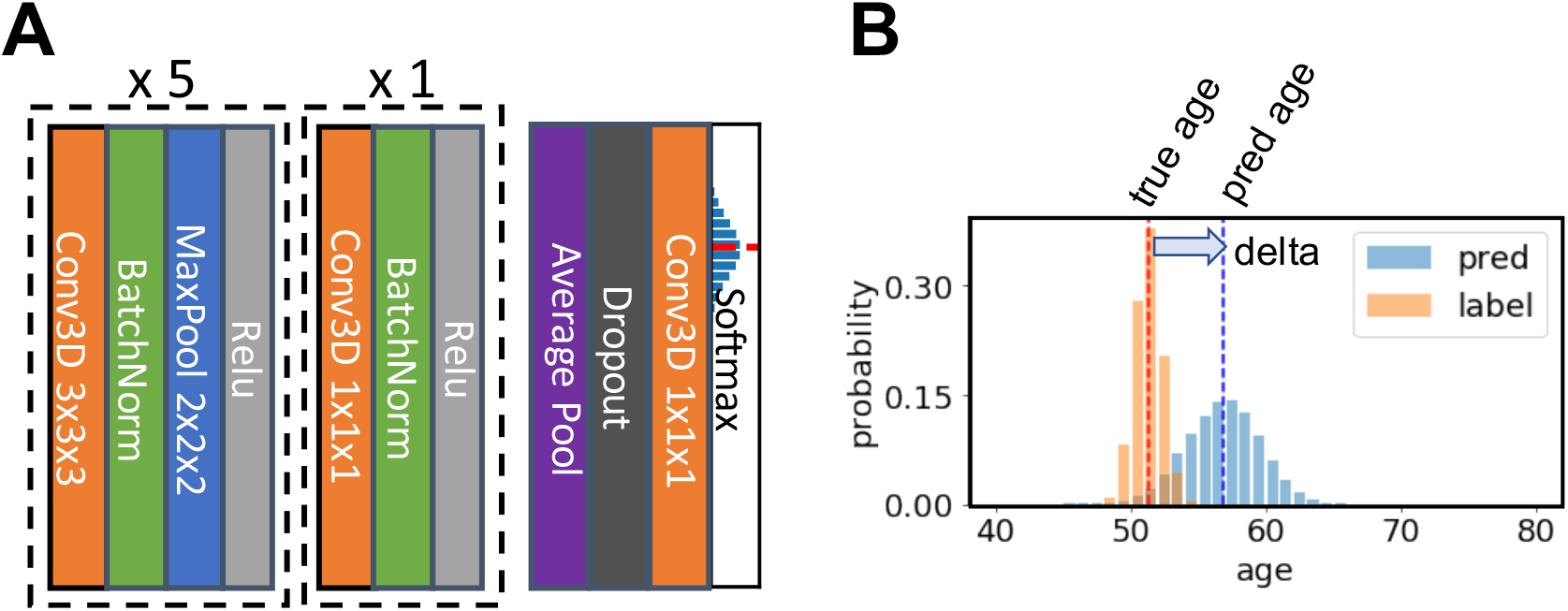
Illustration of the core network for the Simple Fully Convolutional Neural Network (SFCN) model. A) SFCN. The model takes 3D brain image data and contains 7 blocks. Each of the first 5 consecutive blocks consists of a 3×3×3 3D convolution layer, a Batch Norm layer, a Max Pooling layer and a ReLU activation. The 6^th^ block contains one 1×1×1 3D convolution layer, a Batch Norm layer and a ReLU activation. The 7^th^ block contains an average pooling layer, a dropout layer, an 1×1×1 3D convolution layer and a softmax layer. B) An example of soft labels and output probabilities. The soft label is a probability distribution centered around the ground-truth age, and is used to compute the KL-divergence loss, enabling a smooth decrease in the loss function when the prediction improves during the training phase.

The internal process of the model can be interpreted as three stages: 1) The first five blocks extract feature maps from each input image. 2) The sixth block further increases the nonlinearity of the model by adding one extra nonlinear layer but without changing the output size of feature maps. 3) The seventh block maps the generated features to the predicted age probability distribution. The first two stages encode the input image to a feature vector, and the third stage can be viewed as a classifier based on the deep feature. At the first stage, the spatial information is maintained and takes most of the memory. To reduce the overall GPU memory consumption, we limited the channel numbers of the first layer to 32 and put only one convolutional layer in every block. To compare, a VGGnet usually has two convolutional layers inside a block and has 64 channels in the first layer (Simonyan and Zisserman, 2014). At the later stages (higher-level layers) of deep learning models, fully connected (FC) layers usually have the largest number of learnable parameters. For example, the penultimate layer of VGGnet (FC-4096) consists of about 16 million parameters. By removing most of the FC layers and keeping the number of channels small in the last two stages in SFCN, the number of learnable parameters is greatly reduced. Although reducing the number of FC layers can potentially reduce the nonlinearities learnt by the model, in most neuroimaging classification tasks, the number of classes is smaller than that for natural images. For example, there are only 40 “classes” (age bins) for brain age prediction, which is a very small number compared to 1000 classes in the ImageNet classification task. In this case, the small parameter number and the lack of FC layers do not harm the testing (validation) performance.

To compare SFCN with a popular CNN architecture, we implemented a 3D version of ResNet (He et al., 2016). The architecture of 3D-ResNet follows the literature but the convolution filters are changed to 3D. For the experiments, the SFCN and the ResNet share the same training parameters and both achieve successful performance in the training set. (Comparison against a broader set of alternative approaches is provided via the results from the PAC 2019 competition.)

SFCN contains only 3.0 million parameters, which is less than one tenth of the 33.2 million for 3D ResNet-18, 46.2 million for 3D ResNet50 and 133 million for 2D VGGnet (Simonyan and Zisserman, 2014).

### 2.2 Regression models: Elastic Net

We compared our deep learning model with simpler machine learning models using T1 MRI derived features as inputs (Schulz et al., 2019). We choose Elastic Net for our comparison, because it has been shown to be a high-performance and stable machine learning model for neuroimaging data (Jollans et al., 2019). Three forms of the T1 data from UKB were (separately) used for age prediction: (1) Voxel-level linearly registered “raw” T1 images; (2) Voxel-level grey matter partial-volume estimated by FSLVBM voxel-based morphometry; (3) T1-image derived region-level phenotypes (Miller et al., 2016). In the training set, we used principal component analysis (PCA) to reduce the data into an L-dimensional space (L=5000 for (1) and (2), no PCA for (3)), and then used Pearson correlation to select the top k features (from k=10 to all features) that correlate with age, and finally used elastic net regression (implemented in the glmnet package) to predict age (Friedman et al., 2010). All model parameters were optimised via internal cross-validation within the validation set. The selected best model was applied to the test set and performance reported. Besides age prediction, the logistic version of elastic net is also implemented with the scikit-learn package (Pedregosa et al., 2011) for sex classification on the T1-image derived region-level phenotype features. We also implemented a widely-used support vector machine for brain age prediction, but did not find better performance compared with the above model.

### 2.3 Bias correction

We used the linear bias correction method described in (Smith et al., 2019) for bias correction for the delta. Such a bias correction is valuable for most brain-age prediction studies, as there is normally an underfitting of the prediction, due to problems such as regression dilution and non-Gaussian age distribution. Defining *y* to be chronological age and *x* the predicted age, we fitted a linear regression *x* = *ay* + *b* to the left-out validation set (with labels). The corrected predicted age is estimated by

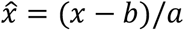

This method requires (at the point of estimating a and b from x and y) that the chronological ages are known. For the label-missing (final evaluation) test set, we assumed that *a* and *b* are generalisable, and used the coefficients previously fitted in the left-out validation set to estimate the corrected brain-age delta.

## 3 Experiments

### 3.1 Datasets and preprocessing

#### 3.1.1 UK Biobank

UK Biobank is collecting a large-cohort of brain imaging data from predominantly healthy participants (Miller et al., 2016). In this study, we used the T1 data from 14,503 subjects (mean age 52.7 years, standard deviation 7.5 years, range 44-80 years), of which 12,949 were used for training, 518 for validation and 1,036 for testing. The image preprocessing pipeline is described in (Alfaro-Almagro et al., 2018). We used data as preprocessed already (by our laboratory on behalf of UK Biobank), and as available to all researchers who have been granted access to UKB data. The input data to the deep neural network model was brain extracted, bias corrected and linearly registered to MNI152 standard space (unless otherwise specified). The head size of subjects is normalized as a result of the linear registration.

#### 3.1.2 PAC 2019

As part of testing the performance of our method objectively, we participated in the Predictive Analytic Challenge (PAC) 2019. This competition was broken down into two parts: a) to achieve the lowest mean absolute error 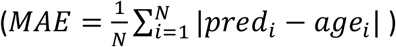 for brain age prediction; and b) to achieve the lowest MAE while keeping the Spearman correlation between the brain-age delta and the chronological age under 0.1 (because in general, ideally delta would have no bias or age dependence). The dataset contains T1 structural MRI brain images from 2,638 subjects (mean age 35.9 years, standard deviation 16.2 years, range 17-90 years).^2^ We used 2,199 subjects for training, and 439 subjects as a left-out validation set. In addition, there were 660 subjects whose labels were unknown to the challenge participants, forming a test set for benchmarking (i.e., the results on this test set determined the final challenge scores).

### 3.2 Training and testing

During the training process, we use a Stochastic Gradient Descent (SGD) optimiser (Sutskever et al., 2013) for the UKB dataset to minimise a Kullback–Leibler divergence loss function between the predicted probability and a Gaussian distribution (the mean is the true age, and the distribution sigma is 1 year for UKB) for each training subject. This soft-classification loss encourages the model to predict age as accurately as possible. To reduce over-fitting, two data augmentation methods are applied during the training phase. In every epoch, the training input is 1) randomly shifted by 0, 1 or 2 voxels along every axis; 2) has a probability of 50% to be mirrored about the sagittal plane.

The performance of the model can be evaluated by Mean Absolute Error (MAE) and Pearson correlation coefficient (r-value) in the validation and test sets.

All the models were trained with two NVIDIA P100 GPUs. The training time was approximately 0.5 hour to go through each of the 12,949 training subjects once (i.e., one training epoch). The L2 weight decay coefficient was 0.001. The batch size was 8. The learning rate for the SGD optimiser was initialized as 0.01, then multiplied by 0.3 every 30 epochs unless otherwise specified. The total epoch number is 130 for the 12,949 training subjects. The epoch number is adjusted accordingly for the experiments with smaller training sets so that the training steps are roughly the same. The epoch with the best validation MAE is used for testing. The deep learning models are trained with the same hyperparameters, as we found that model performance is stable against small changes of hyperparameters.

For the ensemble strategy, we randomly initialised and trained 20 models; 5 (identical network structure but randomly-initialised parameters) models were trained on each of the four input data types: linearly registered GM and WM, non-linearly registered T1 and linearly registered T1. The ensemble experiments use 2,590 subjects for training to reduce the overall computation time. The prediction is made by averaging the results of all 20 models.

### 3.3 Sex classification

To show the generalisability of SFCN to other tasks, we also tested the performance for sex classification. The input brain is linearly registered to standard space (same as the age prediction experiments) so that the overall brain size is the same for all subjects of both sexes. The architecture of the model and the training setting remains mostly the same as for age prediction, with differences now described. With all other model parameters unchanged, the number of classes is set to two and the loss function is changed to cross entropy. The learning rate for the SGD optimiser was initialized as 0.01, then multiplied by 0.3 every 30 epochs. The total epoch number is 220 for the experiments with 100 training subjects, and 150 for all other experiments (1036, 4662, 9841 training subjects, respectively).

## 4 Results

### 4.1 The performance of SFCN in UK Biobank data

Table 1 shows the performance of the SFCN in the UKB dataset with 12,949 training subjects. SFCN with data augmentation and dropout achieved an MAE of 2.14 years, which is 0.46 years better than that without these regularizations.

**Table 1.** Performance of different deep learning models in UK Biobank data. The training set size is 12,949. The input data are T1 MRI images which are linearly registered to a standard space. After the training is done, the epoch with the best validation MAE is selected to be evaluated on the test set. The test results are bootstrapped 1000 times to compute the mean test MAE and the standard deviation. Epochs 95 to 110 are selected to compute the train MAE, standard deviation of the train MAE and the mean validation-train MAE gap (the difference between the validation MAE and the train MAE). All models used the same regularization techniques (dropout, voxel shifting, and mirroring) except for the first row (SFCN without regularization). All models are trained with SGD (except for the second row with ADAM).

| Model | Test MAE (years) | Validation MAE (yrs) | Train MAE (yrs) | Val-Train MAE gap (yrs) |
| --- | --- | --- | --- | --- |
| SFCN without regularization | 2.60±0.06 | 2.62±0.03 | 0.337±0.012 | 2.30±0.03 |
| SFCN (ADAM) | 2.39±0.06 | 2.604±0.009 | 1.610±0.011 | 0.993±0.015 |
| **SFCN (SGD)** | **2.14±0.05** | **2.18±0.04** | **1.36±0.03** | **0.83±0.06** |
| 3D ResNet18 | 2.50±0.06 | 2.38±0.03 | 0.40±0.04 | 1.98±0.05 |
| 3D ResNet50 | 2.32±0.05 | 2.33±0.03 | 0.88±0.05 | 1.45±0.05 |
| 3D ResNet101 | 2.41±0.06 | 2.43±0.02 | 0.85±0.04 | 1.59±0.04 |
| 3D ResNet152 | 2.38±0.06 | 2.38±0.03 | 1.06±0.05 | 1.32±0.06 |

With the same regularization techniques, we then compared SFCN with other popular CNN architectures, namely the 3D version of ResNet18, ResNet50, ResNet101 and ResNet152 (He et al., 2016). Unlike in the ImageNet classification task (He et al., 2016), deeper models do not perform better than the shallow ones in brain age prediction: ResNet50 presents the best MAE among the four models (MAE = 2.32 years), rather than the deeper ones with the same basic units (ResNet101, MAE = 2.41 years; ResNet152, MAE = 2.38 years). Yet, SFCN is 0.18 years better than the best ResNet model.

During the hyperparameter tuning phase, we noted that the choice of optimizer may affect the model performance. To demonstrate this, we train an SFCN model with 12,949 subjects using ADAM optimizer (Kingma and Ba, 2014). The model converges to a less optimal point with test MAE = 2.39 years, which is 0.15 years worse than its SGD counterpart, and has a larger validation-train MAE gap.

Among all the tested architectures, the most lightweight model, SFCN, achieves the best performance. As shown in Table 1, the SFCN with regularizations achieved the best test performance but by far the worst in the training set, suggesting a significant difference (between models) in levels of over-fitting. The gap (a measure of over-fitting) between the validation MAE and the train MAE is 0.83 years for the SFCN, which is the smallest among all the models.

The SFCN model trained with a dropout layer and data augmentation achieves the best MAE. To study the effect of the regularisation techniques, we trained models each with one of the three techniques, namely, dropout, voxel shifting and mirroring, and show the test results in Table 2, with 2,072 training subjects (hence the worse overall results compared with the above). When applied to each of these 3 techniques individually during training, each of the regularisation methods reduces over-fitting and improves the test MAE by about 0.1-0.5 years. Combining all the three methods together, the model achieves the best test performance given this number of training subjects (MAE = 2.82 years), showing a large improvement of 0.85 years compared with the unregularized model. We also experimented replacing BatchNorm layers (Ioffe and Szegedy, 2015) with InstanceNorm layers (Ulyanov et al., 2016), which achieves comparable MAE. Finally, we added one fully connected layer with 64 channels (together with batch normalisation) before the final layer. While giving similar training MAE, the added layer reduces the generalisability to the test set (test MAE=3.59y). Our results clearly show that the regularisation techniques improve the model performance and the lightweight model structure outperforms the tested deep models. These observations can be used for future reference to design deep learning strategy in neuroimage datasets.

**Table 2.** Performance of SFCN with different regularisation and data augmentation methods. DP=Dropout, VS=Voxel Shifting, MR=Mirroring, 1FC = SFCN with one extra fully connected layer. The training set size is 2,072. The input data are T1 MRI images which are linearly registered to a standard space. After the training is done, the epoch with the best validation MAE is selected to be evaluated on the test set. The test results are bootstrapped 1000 times to compute the mean test MAE and the standard deviation. Epochs 185 to 200 are selected to compute the train MAE, standard deviation of the train MAE and the mean validation-train MAE gap (the difference between the validation MAE and the train MAE).

| Data Augmentation and Regularisation | Test MAE (yrs) | Validation MAE (yrs) | Train MAE (yrs) | Val-Train MAE gap (yrs) |
| --- | --- | --- | --- | --- |
| None | 3.67±0.09 | 3.59±0.02 | 0.305±0.007 | 3.29±0.03 |
| DP | 3.59±0.08 | 3.67±0.04 | 0.92±0.02 | 2.76±0.05 |
| VS | 3.05±0.07 | 3.14±0.03 | 0.59±0.01 | 2.56±0.03 |
| MR | 3.13±0.08 | 3.25±0.02 | 0.47±0.01 | 2.78±0.03 |
| DP + VS + MR | **2.82±0.07** | **2.73±0.03** | **1.51±0.03** | **1.22±0.04** |
| DP + VS + MR + 1FC | 3.59±0.08 | 3.6±0.3 | 1.49±0.05 | 2.1±0.3 |
| DP + VS + MR + InstanceNorm | 2.86±0.07 | 2.83±0.02 | 1.70±0.02 | 1.13±0.03 |

Our presented strategy achieves state-of-the-art results in brain age prediction. Table 3 shows a summary of previously reported brain age prediction MAE results (Kolbeinsson et al., 2019; Ning et al., 2018; Smith et al., 2020). To eliminate the effect of sample size differences (i.e., to make these comparisons as meaningful as possible), we trained SFCN with comparable training set sizes as the previous studies, and compared performance with those. With about 2600 training subjects, SFCN achieves an MAE of 2.76 years, while linear regression achieves MAE 3.5 years (Ning et al., 2018). With about 5000 training subjects, SFCN achieves 2.28 years MAE while 3D-ResNet with tensor regression achieves 2.58 years (Kolbeinsson et al., 2019). For the larger training set with more than 10,000 subjects, linear regression with multi-modality IDPs (including fMRI and DTI features) achieves an MAE of 2.9 years (Smith et al., 2020, 2019), whereas SFCN obtains the best MAE in UK Biobank with an MAE of 2.14 years.

**Table 3.** A summary of the reported UK Biobank study in brain age prediction.

| Training set size | Model | Performance |
| --- | --- | --- |
|  |  | MAE (yrs) |
| **2590** | **SFCN** | **2.76±0.06** |
| 2679 | Linear regression Ning et al. 2019 bioRxiv | 3.5 |
| **5180** | **SFCN** | **2.28±0.05** |
| 5700 | 3D-ResNet + Tensor Regression Kolbeinsson et al. 2019 arxiv | 2.58 |
| **12949** | **SFCN** | **2.14±0.05** |
| 18707 | IDP + ICA + Linear regression Smith et al. 2020 eLife | 2.9 |

Besides the state-of-the-art brain age MAE performance among all the reported studies in UK Biobank, our model and strategy also achieved 99.5% accuracy in the hold-out test set for sex classification (0.5% error rate) based on T1 images, as summarized in Table 4. This result is a considerate improvement compared to the previously reported results (classification accuracies varying from 69% to 93%, with or without head size regressed out) (Chekroud et al., 2016; Giudice et al., 2016; Joel et al., 2016, 2015; Rosenblatta, 2016). This result suggests that SFCN is generalisable to other tasks for neuroimaging research.

**Table 4.**
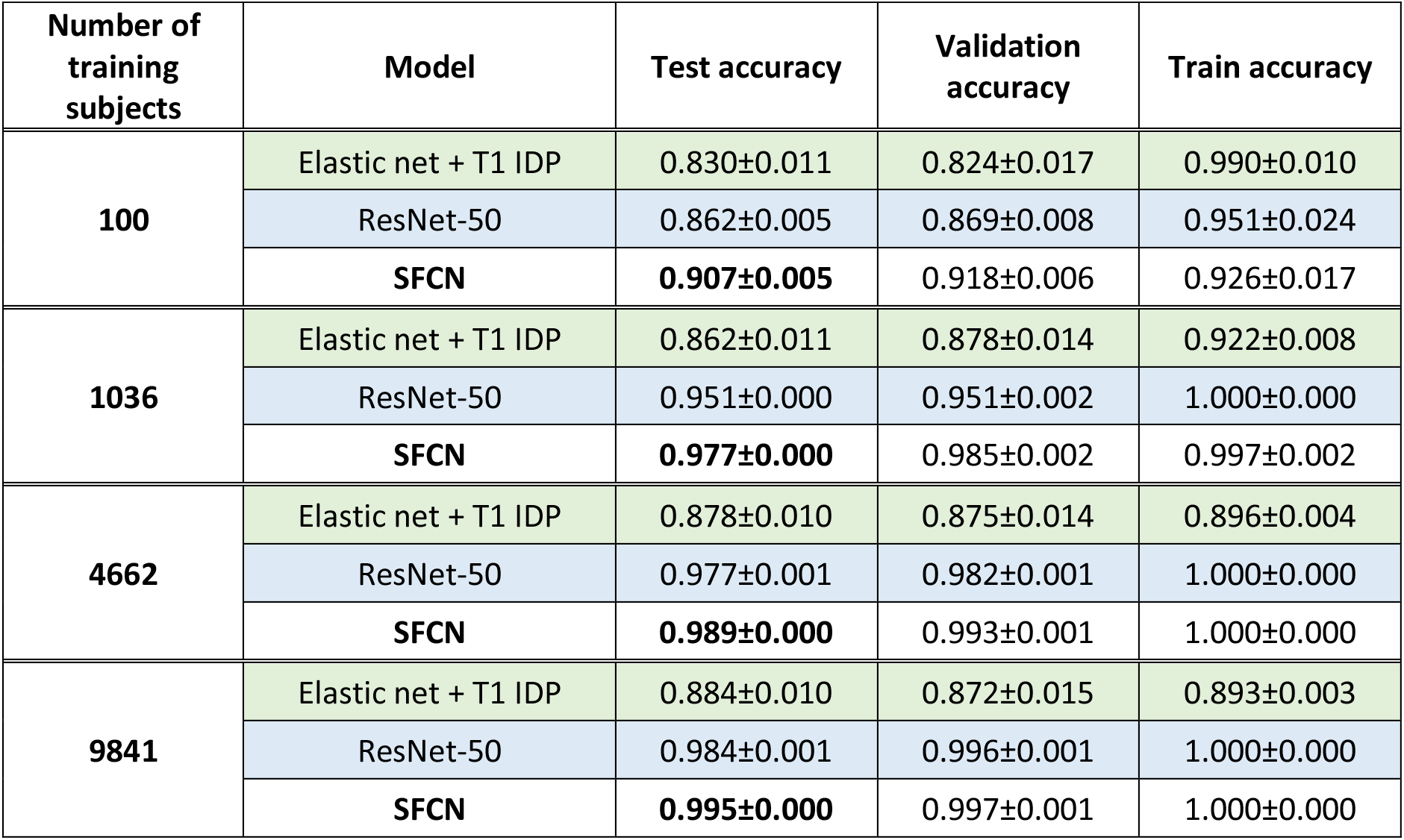
Sex prediction accuracy of different deep learning models in UK Biobank data with different numbers of training subjects. The input data are T1 MRI images which are linearly registered to a standard space. After the training is completed, the final epoch is used to be evaluated on the test set. For SFCN and ResNet-50, last 20 epochs are used to compute the validation/train accuracy and the standard deviations, and every 5 out of the last 20 epochs are used to compute test accuracy. For elastic net, 1000-fold bootstrapping is used to compute the mean and the standard deviation of the test/validation/train accuracy.

| Number of training subjects | Model | Test accuracy | Validation accuracy | Train accuracy |
| --- | --- | --- | --- | --- |
| **100** | Elastic net + T1 IDP | 0.830±0.011 | 0.824±0.017 | 0.990±0.010 |
|  | ResNet-50 | 0.862±0.005 | 0.869±0.008 | 0.951±0.024 |
|  | **SFCN** | **0.907±0.005** | 0.918±0.006 | 0.926±0.017 |
| **1036** | Elastic net + T1 IDP | 0.862±0.011 | 0.878±0.014 | 0.922±0.008 |
|  | ResNet-50 | 0.951±0.000 | 0.951±0.002 | 1.000±0.000 |
|  | **SFCN** | **0.977±0.000** | 0.985±0.002 | 0.997±0.002 |
| **4662** | Elastic net + T1 IDP | 0.878±0.010 | 0.875±0.014 | 0.896±0.004 |
|  | ResNet-50 | 0.977±0.001 | 0.982±0.001 | 1.000±0.000 |
|  | **SFCN** | **0.989±0.000** | 0.993±0.001 | 1.000±0.000 |
| **9841** | Elastic net + T1 IDP | 0.884±0.010 | 0.872±0.015 | 0.893±0.003 |
|  | ResNet-50 | 0.984±0.001 | 0.996±0.001 | 1.000±0.000 |
|  | **SFCN** | **0.995±0.000** | 0.997±0.001 | 1.000±0.000 |

### 4.2 Comparing the learning curves of SFCN with simpler regression models

There are controversies regarding whether a deep learning model can perform better than linear models to predict phenotypic and behavioural variables using neuroimaging data (He et al., 2020; Schulz et al., 2019). For the task brain age prediction using T1 structural MRI data, two questions remain to be answered: 1. Do DL models surpass the performance of simpler regression models? 2. How many training samples do DL or simpler regression models need for good performance?

We compared our deep learning model and a well-tuned regression model, elastic net, for brain age prediction. We also explored the effect of the training dataset size (from 50 to 12,949 subjects) on the performance of the two models. As summarised in Figure 2, we find that the SFCN outperforms elastic net regardless of the training set size. Even with as few as 50 training subjects, the DL model achieves better performance.

**Figure 2.**
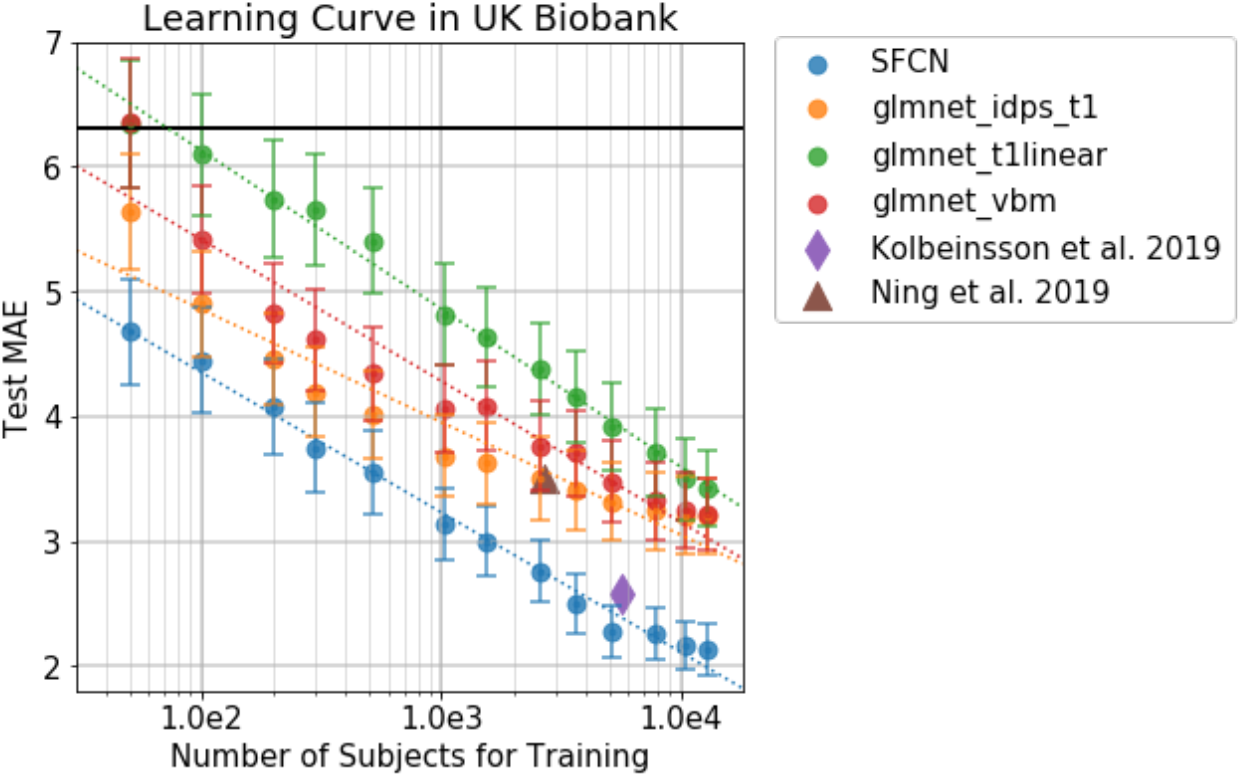
Learning curve for SFCN in UK Biobank data. The methods include SFCN, Elastic nets (glmnet) with different input features, and existing studies. The error bars show the standard deviation by 1000 bootstrap samples. The dashed lines show the log-linear relationship between the training set size and the testing MAE. This shows that as the dataset size doubles, the MAE decrease by around 0.3 to 0.4 years for the linear and the deep learning models. The horizontal dark solid line indicates the MAE when using the population mean age as the predicted age for every testing subject.

The same data-efficiency is also seen in the sex classification task. As summarized in Table 4, in all the four training settings (small dataset: 100 training subjects, medium dataset: 1036 training subjects, large dataset: 4662/9841 training subjects), deep learning methods achieve higher sex classification accuracy than the elastic net.

MAE decreases with the increase of training set size, and approximately follows a linear-log relationship for all methods. If the training set size doubles, the MAE decreases by about 0.3 to 0.4 years. In our experiment setup, there is no conclusive signature of performance saturation for the large dataset size, although the last few data points do deviate from the simple linear-log relationship. With the increasing size of UK Biobank and other datasets, we can expect even better performance in future studies.

### 4.3 Semi-multimodal model ensemble improves the performance with limited number of training subjects

In previous sections, we trained our SFCN model using only one modality, namely raw T1 data linearly registered to the MNI space (Lin). To test whether adding other modalities (here “modalities” refers to different kinds of preprocessing of the T1 data) can further boost performance, we trained SFCN using three other modalities derived from T1 image data: raw

T1 data nonlinearly registered to the MNI space (NonLin), segmented grey matter (GM) and white matter (WM) volumes.

For each of the above 4 modalities, we trained 5 models using different random parameter initializations. To prove the effectiveness of the ensemble strategy without greatly increasing the computing time, we choose a training dataset size of 2,590 subjects for all the models used in this section.

Models trained with different modalities achieve comparable performance with small differences in MAE. NonLin achieved the best MAE (2.73 years), while the Lin and GM achieved comparable MAE of 2.80 years. These modalities are all better than the MAE for WM (2.86 years), as shown in Table 5.

**Table 5.**
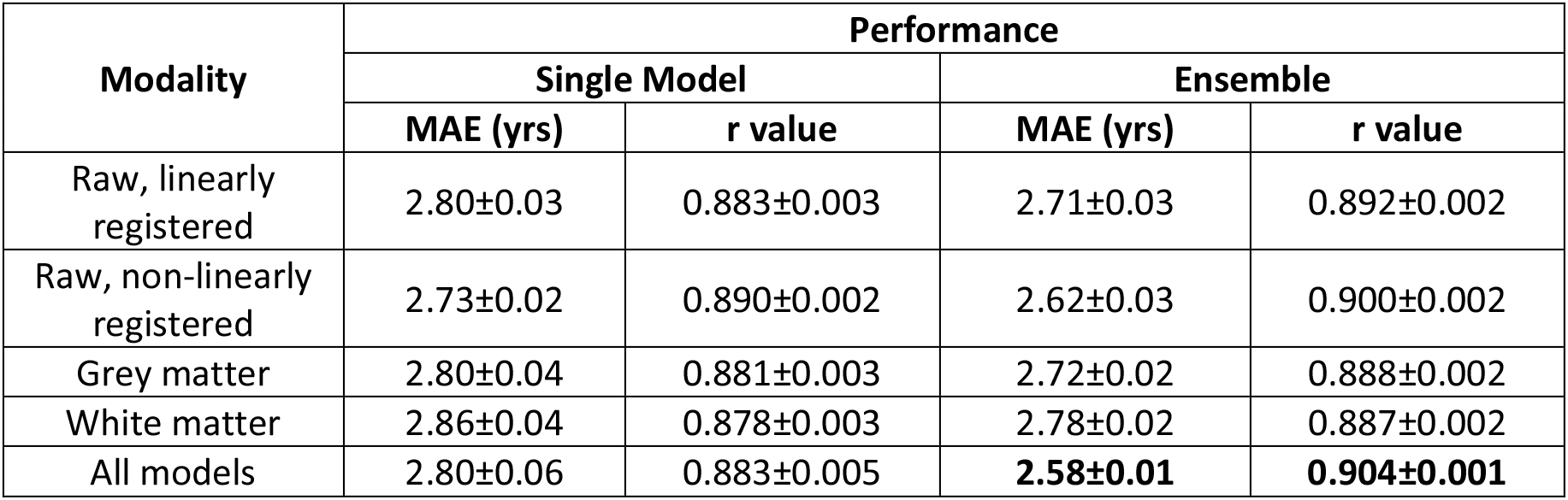
Performance of models trained/tested with different modalities in the test set of the UK Biobank dataset. 5 models were trained for each modality and used to predict brain age individually. The mean and the standard deviation of the single model performances were computed within each modality. For the ensemble performance, 5 models are randomly selected (with duplications allowed) and the predictions were averaged to give the ensemble prediction. This process is repeated for 1000 times to compute the mean test MAE and the standard deviation. For the final ensemble with all modalities, 5 models are randomly selected (with duplications allowed) within each modality and the 20 selected models were used to make the final prediction. This process is repeated for 1000 times to compute the mean test MAE and the standard deviation.

Even though different modalities may result in similar MAEs, the trained models (and deltas) may contain distinct information. This is shown in the correlation matrix of deltas predicted by each of the 20 models in the test sets in Figure 3A (these correlations are between any two estimates of the *N*_*subjects*_ × 1 vector of deltas). Models with the same modalities show higher correlation for the brain-age delta prediction.

**Figure 3.**
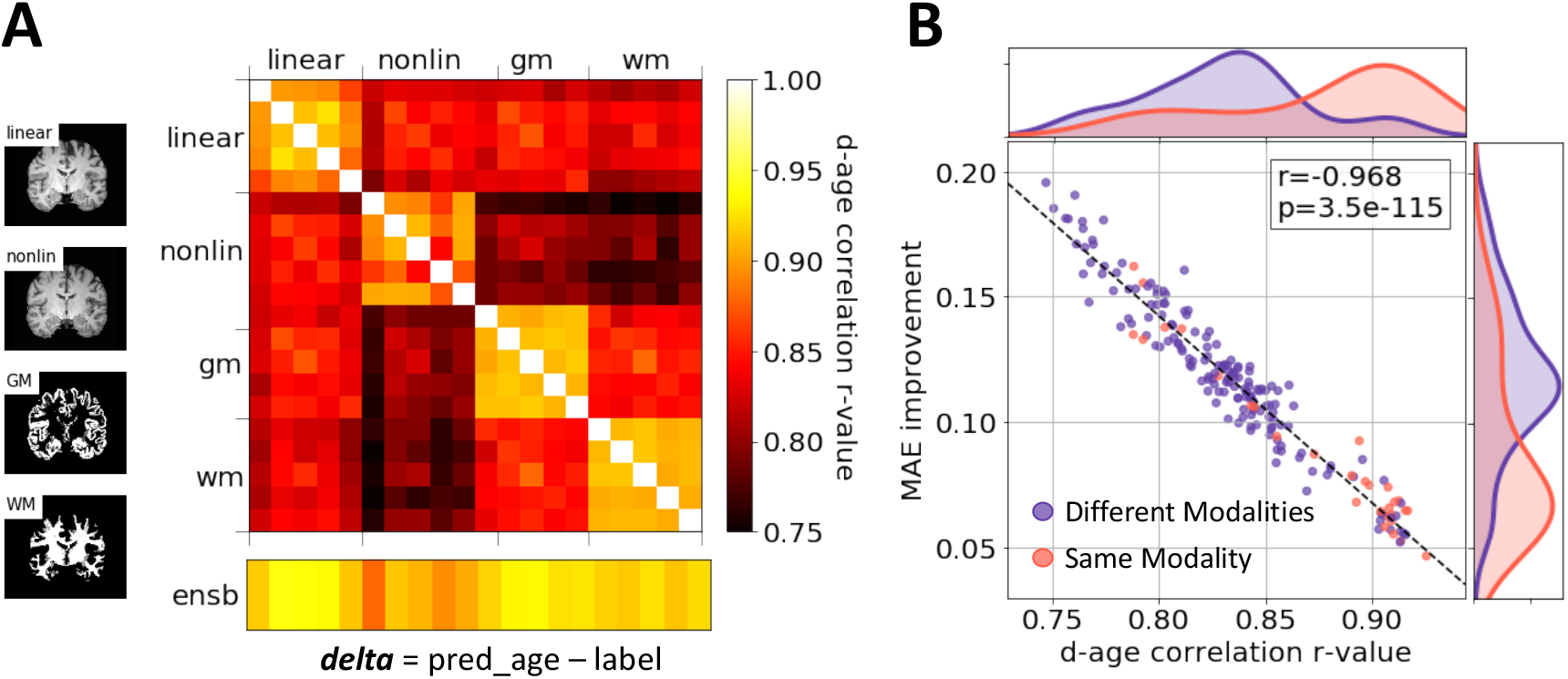
Model ensemble. A) Correlations of brain-age delta predictions between models trained and tested with different modalities. The color coding shows the correlation r-value. For each modality, the training subjects are split into 5 folds, and each model is trained with one-fold being left-out. The delta estimation is made in the common validation set. Any two models trained with the same modality show stronger correlation (between their respective delta estimates) than the models trained with different modalities. The bottom row shows the correlation between individual models and the ensemble prediction. B) Scatter plot: ensemble performance improvement versus delta correlation of any two models. Purple dots represent for ensembles with different modalities. Red dots are for ensembles with the same modality. Normalized histograms of performance improvement and delta correlation of the two-model combinations are plotted alongside. The combination of two models with smaller correlation shows better improvement for the ensemble performance.

To better utilise the information contained within different modalities, we used all four modalities to form an ensemble. For every subject, the 20 models predicted 20 brain ages. The final prediction for the subject was made using the mean of all the predicted ages. This strategy achieved an MAE of 2.58 years, which is 0.22 years better than single model prediction, with 2,590 training subjects.

The success of the ensemble strategy is not only owing to the large number of models, but also due to the independent information gathered from different modalities. To illustrate this, we combined every pair of models and plotted the MAE improvement after ensemble averaging against the delta correlation coefficient in Figure 3B. The result clearly shows that the less correlated two models are, the better performance the ensemble will produce. It has been shown that models trained from different modalities tend to be less correlated. Therefore, combining models from different modalities with complementary information gives the greatest performance enhancement.

### 4.4 Bias correction

The next challenge is bias correction. We illustrate the age prediction results of SFCN trained with 12,949 subjects in Figure 4. The predictions tend to bias towards the mean age of the cohort, which means that younger subjects will be predicted to be older and vice versa. This is due to regression dilution (MacMahon et al., 1990) and other factors such as model regularization and non-Gaussian distribution of the labels (true ages) (Smith et al., 2019), and results in a high correlation between brain-age delta (prediction – age) and chronological age (Spearman’s r=-0.39). We followed (Smith et al., 2019) to regress age out of the delta. In the PAC 2019 competition framework, we do not know the label of the test set. In this case, we regressed out age in the 518-subject validation set and then used the estimated bias correction regression coefficients for bias removal in the test set. This process assumes that the bias distribution is the same for both the validation and the test set, and does not require any knowledge of the age labels in the test set. As summarized in Table 6, this technique reduced the bias Spearman’s r-value from −0.37 to 0.03, with an increase of just 0.15 years in the MAE for the validation set. The generalised strategy (for unlabelled data) reduced the r-value from −0.39 to 0.01, with a small increase (0.11 years) in the MAE for the test set.

**Table 6.** Performance of the ensemble model with and without bias correction. The UK Biobank validation set is used to estimate the slope and intercept from a linear fitting, which is then used to generalized in the unseen UK Biobank test set. This strategy, together with SFCN and the ensemble, was used to take first place in the PAC 2019 Brain Age Prediction Challenge.

| Dataset | Performance |  | Performance with Bias Correction |  |
| --- | --- | --- | --- | --- |
|  | MAE (years) | Spearman Correlation delta vs age | MAE (years) | Spearman Correlation delta vs age |
| UK Biobank validation set | 2.10 | -0.37 | 2.25 | 0.03 |
| UK Biobank test set | 2.14 | -0.39 | 2.25 | 0.01 |
| PAC 2019 Brain Age Prediction Challenge | 2.90 | -0.39 | 2.95 | -0.03 |

**Figure 4.**
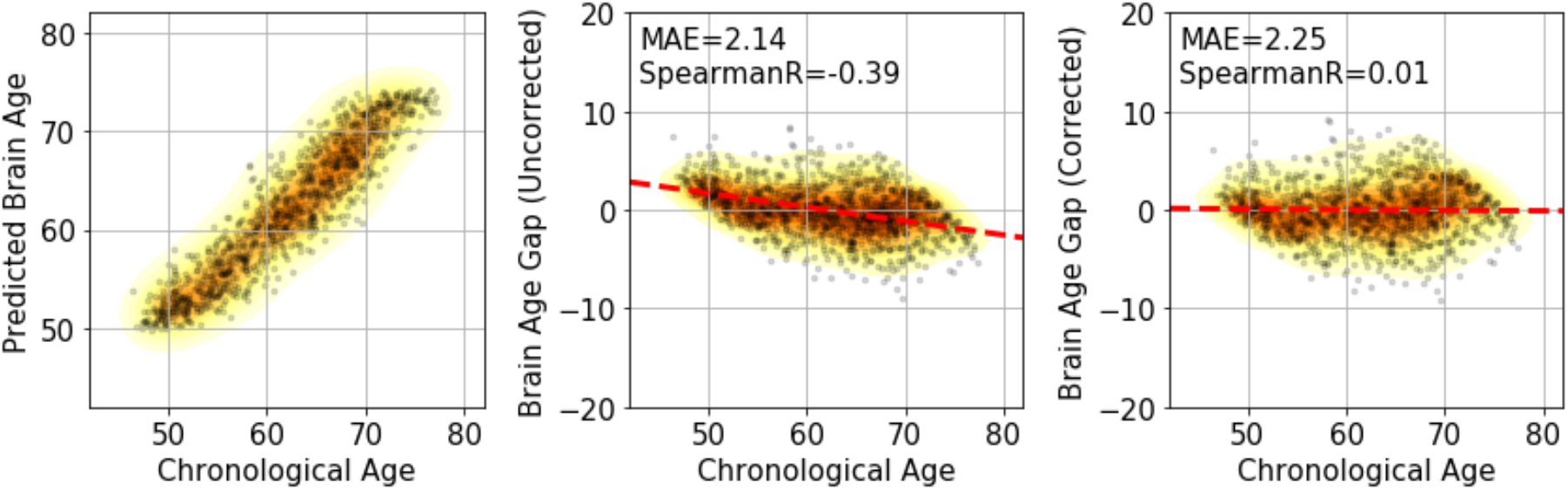
Bias correction. (Left panel) Results of brain age prediction for the UK Biobank test set, SFCN trained with the full training set. (Middle) Results of delta without correction. (Right) Results of delta with correction.

Finally, we tested our methods of SFCN, data augmentation, ensemble and bias correction in the PAC 2019 brain age prediction challenge and achieved first places in both goals of the challenge: 1. to achieve the smallest MAE and 2. To achieve the smallest MAE with bias Spearman’s r-value under 0.1. For the first objective, we achieved MAE=2.90 years, which was 0.18/0.42 years better than the second/third places. For the second objective, we achieved MAE=2.95 years, and this result was 0.85/0.97 years better than the second/third places, i.e., a significant improvement (over the other best approaches) of almost one year. These results are available in the challenge website^3^.

## 5 Discussion

We have demonstrated that with a well-trained model, deep learning method can achieve better performance than the simpler regression models tested even in a training set size as small as 50 subjects. While a few studies have shown the opposite conclusion (that linear regression outperforms deep learning in neuroimaging data (Schulz et al., 2019), we argue that DL method is a large family of algorithms and techniques, some more suitable than others for neuroimaging. Different choices of model architectures and training strategies can result in very different results.

Compared to the simpler regularized regression methods, one of the potential limitations of our method is the training resource consumption. Although the inference time is about a few milliseconds per subject once the model is trained, the training time takes more than 50 hours using 2*P100 GPUs with 14K subjects for one single model and one single modality. As the UK Biobank dataset is still expanding and even larger datasets may appear in the future, the resource requirement will increase as well, and this may limit the ability of fast explorational research. Thus, it is important to develop fast training strategies and/or to test the feasibility of transfer learning to reuse pretrained weights for deep learning models in neuroscience research.

In our study, we have successfully demonstrated the effectiveness of the lightweight model SFCN as an example of a DL method. We have shown that the lightweight architecture without a fully connected layer achieves less overfitting and better results than deeper models in brain age and sex prediction tasks. This observation is in line with the recent study by Raghu et al. (Raghu et al., 2019) which shows that, while deep ImageNet architectures achieve state-of-the-art performance in natural images, small models can achieve comparable (if not better) performance than deeper nets in 2D retina images and chest X-ray images. It is intriguing why the lightweight model performs better than the deeper ones. One insight is that in medical imaging (and neuroimaging) applications, the classification tasks involve only a few classes and thus there is no need for a wide FC layer or large over-parameterization (Raghu et al., 2019). However, it remains an open question how the representations learnt by the lightweight models differ from the deep ones, and what is the general principle to optimally design a minimum architecture for medical imaging applications.

We have also demonstrated the performance gain by including different ‘modalities’ in the model ensemble, rather than using a single modality only. Due to the limitation of the computing power, we use the simplest method (averaging the predictions) and recognise there are more effective ways to combine the information from multi-modality inputs. For example, both Cole et al. (Cole et al., 2017) and Jonsson et al. (Jonsson et al., 2019) trained models with different modalities. Cole et al. concatenated the encodings of different modalities in the FC layer, and Jonsson et al. used a majority voting strategy to form the final prediction, and both received performance gain through multimodal inputs.

We have extended the bias correction method by Smith et al. (Smith et al., 2019) to be able to correct bias in new data where the age is not known, and successfully removed Spearman’s rank correlation between the brain age delta and the true age. However, simple linear regression does not remove nonlinear bias effects. Polynomial regression could be used where supported by (and required by) the data in question, as proposed in (Smith et al., 2019), and using the same extension to new data described in our work.

To conclude, we proposed SFCN, a lightweight deep neural network architecture, which achieved state-of-the-art brain age prediction and sex classification using T1-weighted structural MRI images. We investigated different approaches for boosting the performance of the deep learning model, and tested three factors that are valuable for improving the performance of a single deep learning model in a neuroimaging dataset through a series of controlled experiments: (1) the lightweight model structure (summarised in Table 1), (2) data augmentation and regularisation techniques (e.g., dropout, voxel shifting and mirroring, as summarised in Table 2), (3) large dataset size (summarised in Figure 2). For semi-multimodal data (i.e., data from a single modality but which has been through several distinct processing steps), we presented an ensemble strategy that improved single modality results by utilising the (somewhat) independent information from different modalities. Finally, we showed that regressing the true age out of brain-age delta (predicted age minus actual age) can effectively correct bias, and the fitted slope and intercept can be directly transferred to the unknown test set in UK Biobank, one of the largest neuroimaging datasets. We also showed that, SFCN can outperform simpler regression models even with small training set sizes. These results are successful explorations of the application of DL in the neuroimaging data.

## 6 Acknowledgements

We are grateful for funding from the Wellcome Trust. The Wellcome Centre for Integrative Neuroimaging (WIN FMRIB) is supported by core funding from the Wellcome Trust (203139/Z/16/Z). Funding also from Wellcome Trust Collaborative Award 215573/Z/19/Z. This project is supported by the DeepMedicine project in the Oxford Martin School and the Innovative Medicines Initiative 2 Joint Undertaking under grant agreement No 777394 (for AIMS-2-TRIALS) which receives support from the European Union’s Horizon 2020 research and innovation programme and EFPIA and AUTISM SPEAKS, Autistica, SFARI. This research has been conducted in part using the UK Biobank Resource under Application 8107. We are grateful to UK Biobank for making the data available, and to all UK Biobank study participants, who generously donated their time to make this resource possible. Computation used the Oxford Biomedical Research Computing (BMRC) facility, a joint development between the Wellcome Centre for Human Genetics and the Big Data Institute supported by Health Data Research UK and the NIHR Oxford Biomedical Research Centre.

## Data Availability Statement

The UK Biobank dataset is accessible upon applications via the website: https://www.ukbiobank.ac.uk/. The PAC 2019 dataset consists of several public available datasets and a few datasets provided by the organiser. Interested researchers can apply for the access to the public available datasets as specified in (Cole et al., 2017) and need to contact the PAC 2019 organisers for the rest of the sites.

## Footnotes

1 PAC 2019 website archive: https://web.archive.org/web/20200214101600/https://www.photon-ai.com/pac2019

2 This information is publicly available in the challenge website: https://web.archive.org/web/20200214101600/https://www.photon-ai.com/pac2019

3 Link to the results: https://web.archive.org/web/20200214101600/https://www.photon-ai.com/pac2019#results Team: BrainAgeDifference

